# micronuclAI: Automated quantification of micronuclei for assessment of chromosomal instability

**DOI:** 10.1101/2024.05.24.595722

**Authors:** Miguel A. Ibarra-Arellano, Lindsay A. Caprio, Aroj Hada, Niklas Stotzem, Luke Cai, Shivem Shah, Johannes C. Melms, Florian Wünneman, Benjamin Izar, Denis Schapiro

## Abstract

Chromosomal instability (CIN) is a hallmark of cancer that drives metastasis, immune evasion and treatment resistance. CIN results from chromosome mis-segregation events during anaphase, as excessive chromatin is packaged in micronuclei (MN), that can be enumerated to quantify CIN. Despite recent advancements in automation through computer vision and machine learning, the assessment of CIN remains a predominantly manual and time-consuming task, thus hampering important work in the field. Here, we present *micronuclAI*, a novel pipeline for automated and reliable quantification of MN of varying size, morphology and location from DNA-only stained images. In *micronucleAI*, single-cell crops are extracted from high-resolution microscopy images with the help of segmentation masks, which are then used to train a convolutional neural network (CNN) to output the number of MN associated with each cell. The pipeline was evaluated against manual single-cell level counts by experts and against routinely used MN ratio within the complete image. The classifier was able to achieve a weighted F1 score of 0.937 on the test dataset and the complete pipeline can achieve close to human-level performance on various datasets derived from multiple human and murine cancer cell lines. The pipeline achieved a root-mean-square deviation (RMSE) value of 0.0041, an R^2^ of 0.87 and a Pearson’s correlation of 0.938 on images obtained at 10X magnification. We tested the approach in otherwise isogenic cell lines in which we genetically dialed up or down CIN rates, and also on a publicly available image data set (obtained at 100X) and achieved an RMSE value of 0.0159, an R^2^ of 0.90, and a Pearson’s correlation of 0.951. Given the increasing interest in developing therapies for CIN-driven cancers, this method provides an important, scalable, and rapid approach to quantifying CIN on routinely obtained images. We release a GUI-implementation for easy access and utilization of the pipeline.

## Main

Chromosomal instability (CIN) is a hallmark of aggressive cancers ^1–3^, and developmental and age-related disorders ^4^. In cancer, CIN drives tumor progression, heterogeneity, immune evasion, and treatment resistance across a range of tumor lineages ^3,5–10^. CIN may arise from different mutagens (e.g., radiation, mitotic toxins), defects in DNA repair, and most frequently, from errors in chromosome segregation during anaphase ^11–13^. Following asymmetric distribution of chromatin, cells receiving excessive material frequently package DNA in so-called micronuclei (MN). MN are variable in size (ranging from small (0.5-1 µM) to large (10-15µm)), however less than 1/3 of the size of the primary nucleus, may exist in different morphologies and locations in relation to the nucleus, and lack the normal nuclear envelope, which results in frequent rupture and release of double-stranded DNA (dsDNA) to the cytosol ^14,15^. This in turn triggers the cytosolic DNA sensing machinery via cGAS-STING, that may result in production of pro-immunogenic cytokines (e.g., type I interferons) when stimulated briefly, but suppresses cytokine production and promotes STING-dependent pro-metastatic pathways when activated tonically, such as in the case of most CIN-driven cancers ^7,16,17^. Thus, enumerating MN of varying qualities is a useful approach for quantifying CIN and has important functional implications for tumor behavior, immunity responses, and treatment responses.

Several approaches, such as cytogenetic, quantitative imaging, single-cell genomics, etc. can be used for the assessment of CIN ^18^. Among them, quantitative imaging microscopy is widely used for its simplicity, low-cost, and scalability. Here, assessment of CIN is performed through the quantification of its associated structural biomarkers, including MN, anaphase bridges, and Nuclear Buds (NBUDs), which are MN that may maintain a visible stalk of nucleoplasmic material ^19^, among others ^12,20,21,22^. Although quantification of these functional CIN surrogates from microscopy images is a routine process in cancer research, it is typically achieved through manual scoring. Therefore, microscopy images are divided into high power fields of view (FOV), and the total number of nuclei as well as the number of MN are counted within each FOV. The rate of MN and nuclei is then estimated over the whole image in terms of ratio between the number of MN and the number of nuclei averaged over all the FOV regions. During a standard analysis, multiple images are scored per sample, resulting often in at least 30 FOVs consisting of approximately 1000-1500 primary nuclei. Subsequently, all these images must be manually counted and analyzed (Supplementary Fig. 1) ^7,23–25^.

The manual nature of this workflow makes it tedious, time-consuming and error prone. The sheer amount of data makes it impractical for an annotator to accurately survey an entire image, thus retreating to heuristic approaches. Moreover, the morphology and localization of MN post another source of misclassification errors. The complexity of the task is further exacerbated by the unavailability of a standard protocol leading to inter-observer variability while counting. Additionally, density of nuclei in each image and resulting crowding makes it challenging to confidently separate and accurately assign MN events to nuclei for both humans and automated methods ^26^. Thus, improved methods for automated CIN quantification are necessary to increase speed, accuracy and robustness of CIN-related research.

Here, we present *micronuclAI*, a novel deep-learning-based pipeline for assessment of CIN through automated quantification of MN from nuclei-stained images. micronuclAI distinguishes itself over previous methods ^26–32^ as it 1) can quantify for both MN and NBUDs, 2) requires only nuclear staining, 3) is able to work with 10x to 100x image objectives, 4) can work with any segmentation mask, and most importantly 5) is extensively evaluated in multiple cell lines and thus, ready for use by the community (Fig. 1 and Table 1).

**Figure 1:**
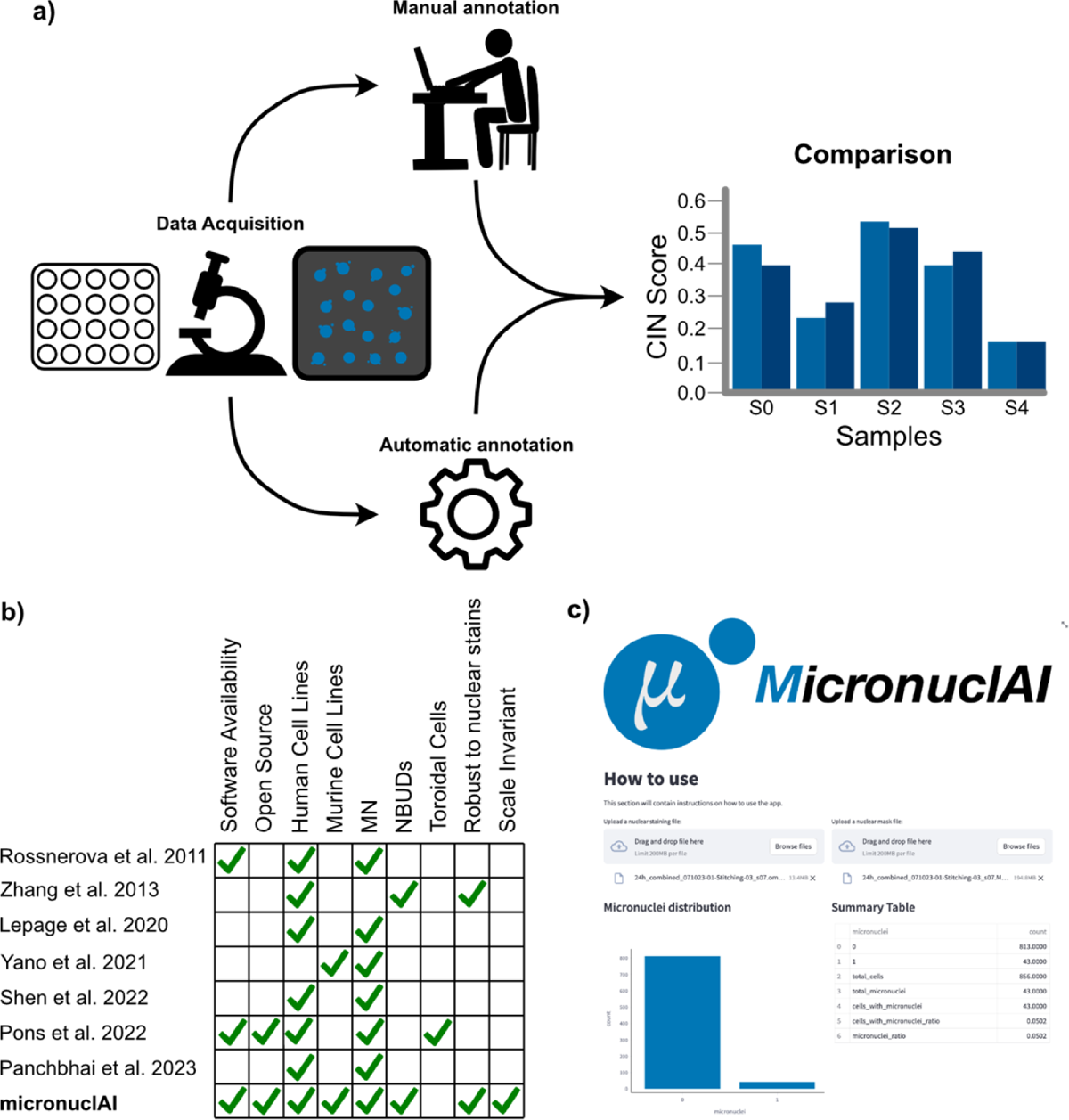
micronuclAI pipeline overview and previous work. **a,** micronuclAI aims to be the go-to tool for CIN assessment via automatic quantification of MN in nuclei-stained images over the manual workflow. Evaluation of the pipeline was performed by comparing the CIN score, a ratio of the total number of micronuclei and associated structures to the total number of nuclei present, both manually and automatically in multiple cell-lines. **b,** Comparison between previous methods and micronuclAI. Prior efforts to automatically quantify CIN bring their own advantages but also have certain limitations involving ability to account for multiple species, multiple MN structures (MN or NBUDs), limited software availability, and dependence on image acquisition parameters. **c,** screenshot of the micronuclAI web application.

**Table 1:**
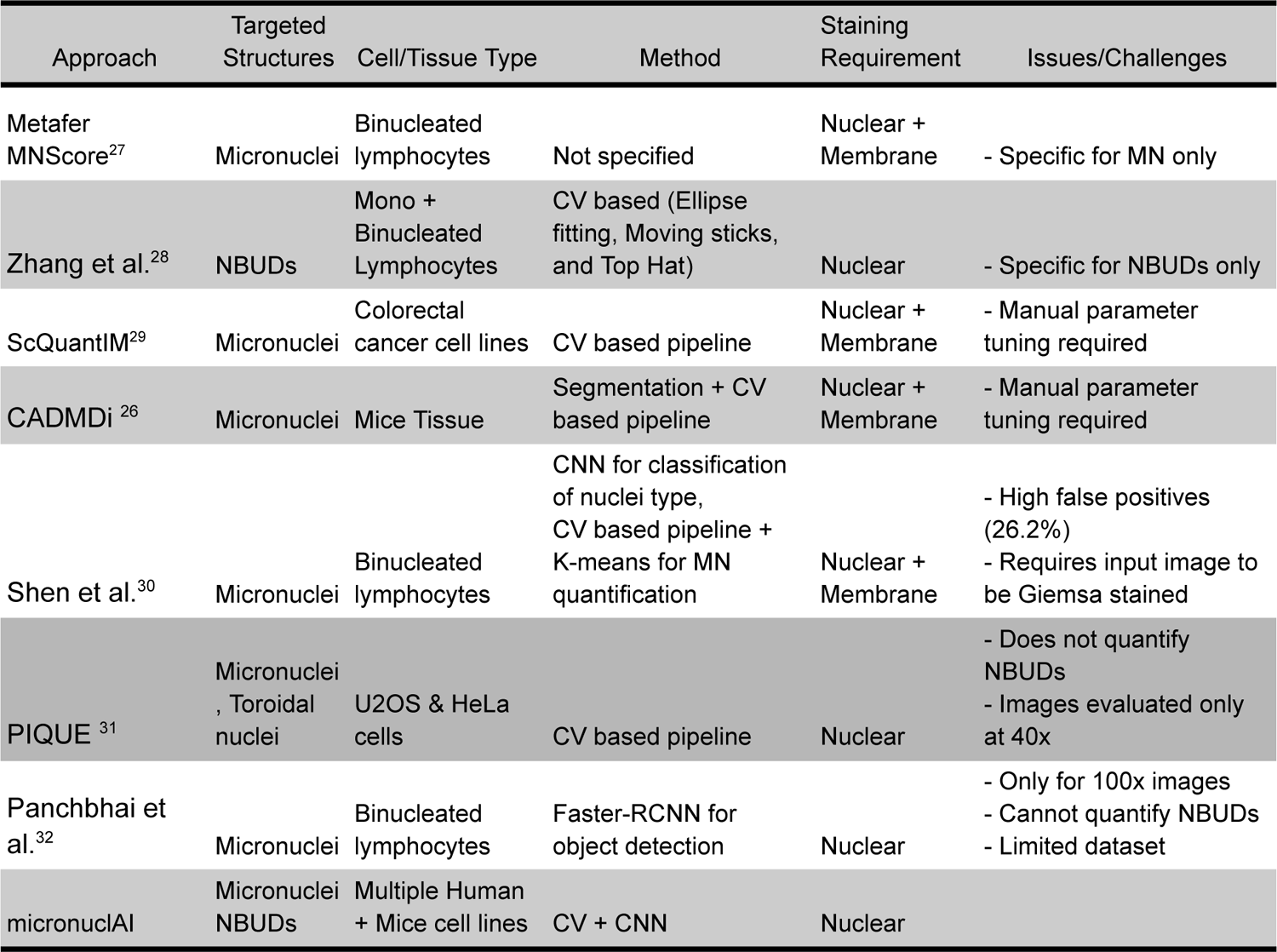
Relevant work.

## Results

### Generation of the training data set

Expert annotation (the process of manually defining the amount of MN) was performed using a custom tool, written in Python ^33^, on 21 images with a resolution of 8829×9072 pixels of the melanoma cell line A375. To perform the nuclear segmentation of the training images, we tested multiple methods and models: DeepCell Mesmer / DeepCell nuclear ^34^, Cellpose ^35^, and Stardist^36^. We visually inspected the generated nuclear segmentation masks and chose the method that generated the most accurate results.

Based on visual inspection, nuclear segmentation was performed using the Stardist segmentation method with default parameters. Importantly, micronuclAI performance is robust across multiple segmentation approaches (Supplementary Fig. 2), thus the choice of segmentation method is not critical. With the help of the segmentation masks, we isolated 84,286 nuclei. From each isolated nucleus, the number of MN present was manually counted for CIN estimation and henceforth termed as “CIN count” (Fig. 2a) (See Methods). In all analyses, NBUDs are considered MN of distinct localization and morphology. From the labeled nuclei, 77,733 (92.23%) had a CIN count of 0; while 6,553 (7.77%) of the labeled nuclei had a CIN count of at least 1. To better handle this data imbalance, we removed 23 (0.0272%) outliers with a CIN count >=4, and randomly sampled the same number of nuclei with a CIN count of 0 to the nuclei with a CIN count >0. The balanced dataset consisted of 13,060 nuclei, from which 6,530 (50%) had CIN count of 0; 5,583 (42.74%) had a CIN count of 1; 597 (4.57%) had a CIN count of 2; and 100 (0.76%) had CIN count of 3 (Fig. 2b). From this balanced dataset, 11,754 nuclei (90%) were used for training-validation and the remaining 1,306 nuclei (10%) as a holdout test dataset (Fig. 3a).

**Figure 2:**
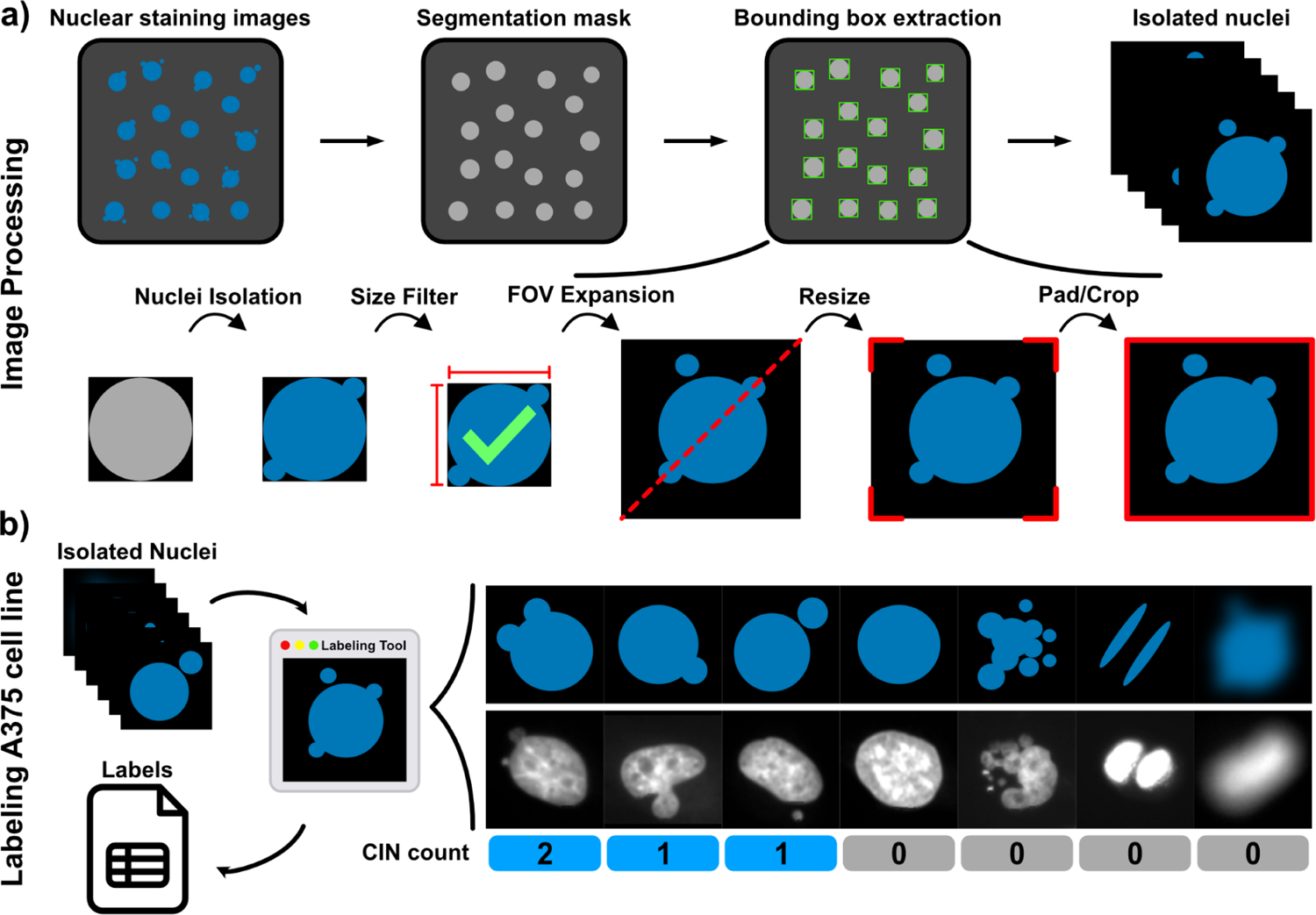
Data preprocessing and labeling. **a,** micronuclAI image preprocessing initially requires the extraction of bounding boxes and isolated nuclei using a segmentation mask. The isolated nuclei are subjected to various preprocessing steps to capture the immediate surrounding and make them homogeneous in size. **b,** With the help of a custom-made labeling tool, the CIN count of each isolated nuclei is recorded. The CIN count quantifies each MN and NBUD per isolated nuclei while specific patterns such as apoptotic nuclei, mitotic nuclei, and low quality or blurry images are given a CIN score of 0 to account for such structures present in real data.

**Figure 3:**
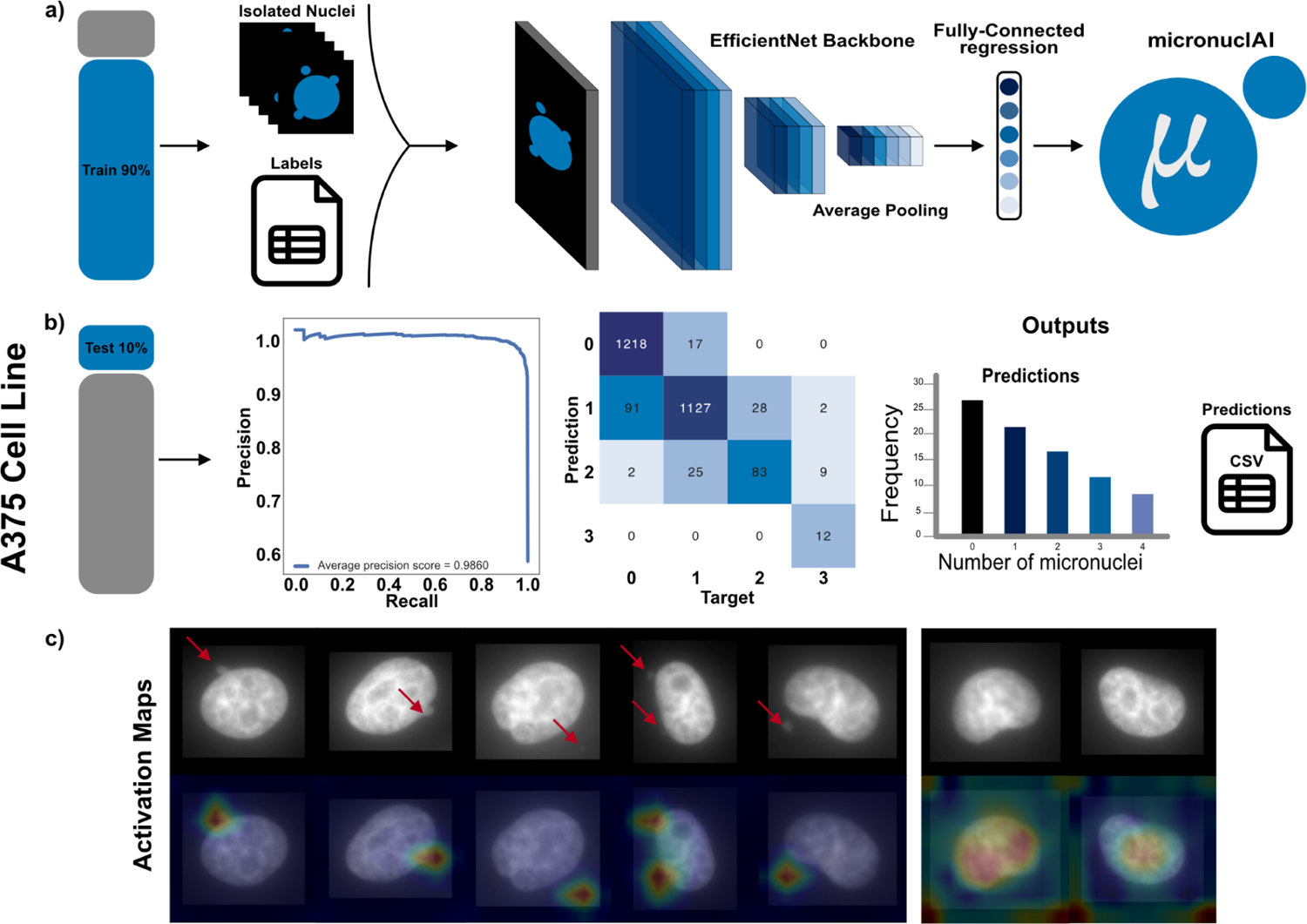
Overview of the model training and testing. **a,** We train the micronuclAI model on the preprocessed and labeled isolated nuclei from DAPI stained images obtained from the A375 cell line. 90% of the labeled data was used for training with a 90:10 train-validation split. We selected the best model based on the performance on the validation set. **b,** The best model was then tested on a holdout test dataset made of the remaining 10% labeled data. With an average precision score of 0.9860 micronuclAI offers high accuracy for the detection of CIN. During inference, micronuclAI outputs predicted CIN counts for each detected nuclei in a csv file along with significant statistics. **c,** Activation maps show localized activation around MN and NBUDs in contrast to nuclei without CIN, where the activation is not localized and appears to diffuse around the primary nuclei.

### micronuclAI accurately identifies CIN in the hold-out test set

On the hold-out test set comprising 1,306 isolated nuclei, the best model (EfficinentNet B0) achieved a high F1-weighted score of 0.9417, and a MCC (Mathew’s correlation coefficient) of 0.8945 (Fig. 3b). These metrics showcase the ability of the model to accurately quantify the number of MN in each isolated nuclei image. Additionally, the attention maps correlate with the areas where MN are present, and in cases where there is no MN, the attention is focused across the entire nucleus (Fig 3c). This shows the model has learned to identify the specific features and patterns associated with MN and conversely accurately identifies cells without MN.

### micronuclAI performance generalizes across biological and technical contexts

To assess the generalizability of micronuclAI to a new cell-line, we evaluated the complete micronuclAI pipeline on images taken from the NCI-H358 non-small cell lung cancer cell lines. The rate of CIN was up- or down-modulated using previously established genetic constructs. For example, overexpression of a dominant negative mutant of mitotic-centromere associated kinesin (dnMCAK) enhances CIN in otherwise CIN-low cell lines ^37^. Thus, in addition to the parental cell line (H358), expression of a vector control (H358-GFP), we also examined CIN-modified derivatives of this cell line (H358-dnMCAK).

Evaluation of the complete pipeline was done based on the calculation of the CIN score, as a ratio of the CIN counts to the number of total nuclei present, compared between manual counts from experts and the automatic counts obtained from micronuclAI. The CIN score for individual images were found to be very accurate for each sample resulting in an RMSE value of 0.0041, R^2^ of 0.881, and a Pearson’s correlation of 0.932. These metrics signify that the manual and automatic counts show a high correlation. Furthermore, the metrics improve when the CIN score is calculated as an average of multiple technical or biological replicate images, which is the standard procedure in manual counting. When taking average counts for the three genetic variants of H358 cell line; H358 control, H358-GFP, and H358-dnMCAK, the values for RMSE decreased to 0.0020, the R^2^ increased to 0.9794, and the Pearson’s correlation increased to 0.989 (Fig. 4a).

**Figure 4:**
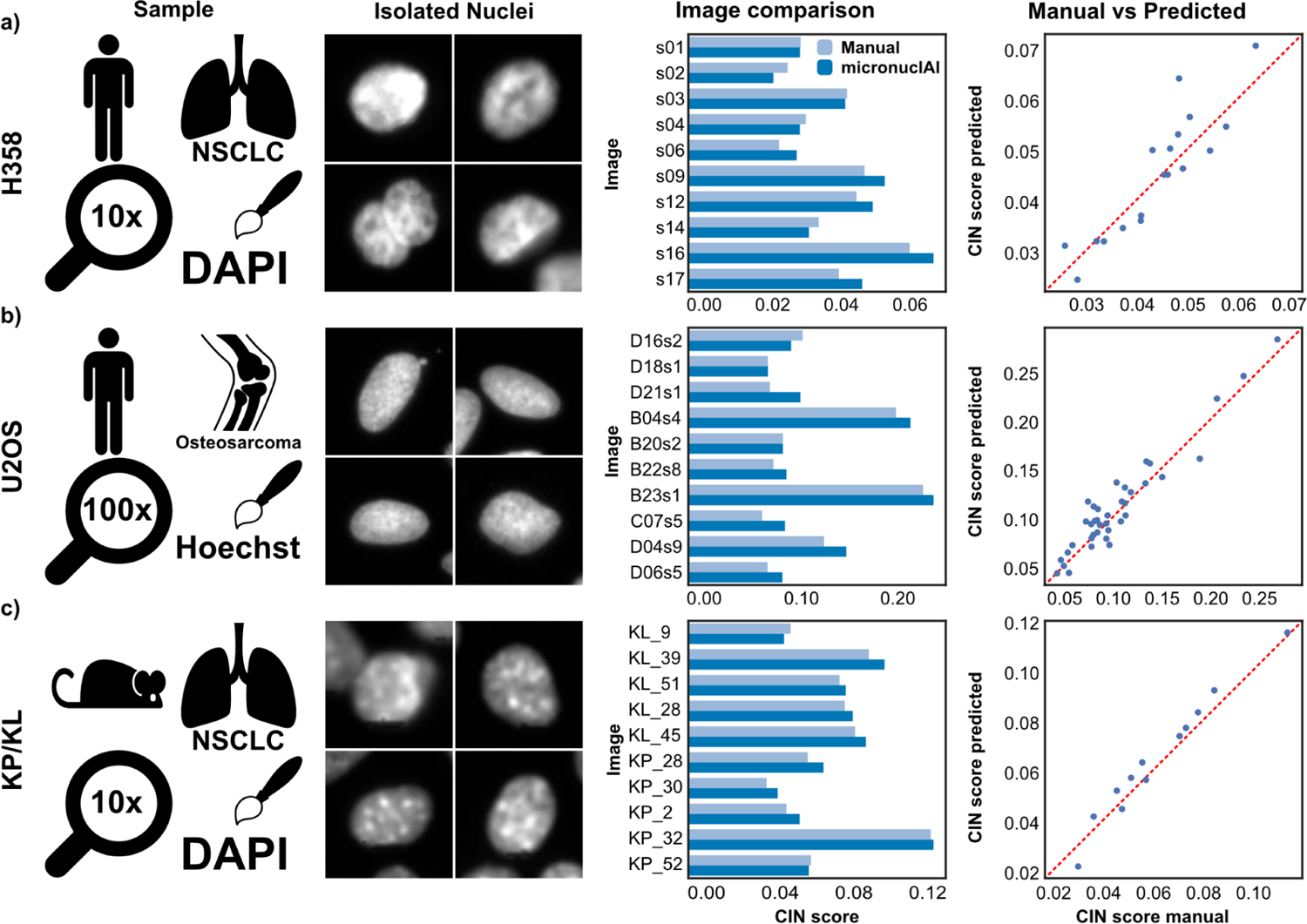
micronuclAI pipeline validation results. To validate the pipeline with a real-world application, we compared manual CIN scores against predictions obtained from micronuclAI for 4 different cell lines: **a,** To test the scalability of our method, we used a DAPI-stained H358 human non-small-cell lung cancer derived cell line (R^2^=0.869, Person’s Corr=0.932). **b,** To test the scale and robustness of the pipeline, we used Hoechst stained U20S human osteosarcoma derived cell line obtained from the Broad Bioimage Benchmark Collection (R^2^=0.905, Pearson’s Corr=0.951). **c,** Lastly, we tested micronuclei’s capability to identify CIN on non-human cell lines by using the KP and KL cell lines derived from mouse non-small-cell lung cancer (R^2^=0.9662, Pearson’s Corr=0.982). For each cell line, we can observe the output of the nuclei isolation step resulting in homogeneous patches, which are then used as input for the model to make predictions on.

### micronuclAI performance is robust to different species, scales and technical variation

We tested micronuclAI’s generalizability to unseen nuclear stains and imaging conditions using an independent dataset, the BBBC039v1 ^38^ image set which are Hoechst-stained U2OS osteosarcoma cell line imaged at 100X. Comparing the CIN score between human expert count and our pipeline revealed that micronuclAI was able to accurately quantify the number of cells and micronuclei in the images. For 20 randomly selected images present in the dataset, micronuclAI was able to achieve an RMSE of 0.0159, an R^2^ of 0.905, and a Pearson’s correlation of 0.951 confirming the scale invariability in images ranging from 10x to 100x scale and stain invariability between DAPI and Hoechst stains (Fig. 4b).

Finally, we evaluated the performance of the pipeline on murine lung cancer cell lines KP (*Kras^G12C/WT^Tp53*^-/-^) (n=5) and KL (*Kras^G12C/WT^Lkb1*^-/-^) (n=5). We observed a great degree of correlation between the results with an RMSE of 0.00536, an R^2^ of 0.9662, and a Pearson’s correlation coefficient of 0.9829. Thus, confirming the robustness of the pipeline to non-human cell lines. Importantly, we used our previously identified default parameters for preprocessing and predictions of these cell lines, thus, further optimization would likely improve the results (Fig. 4c). All the results obtained on the different cell types have been summarized in Table 2.

**Table 2:** Results obtained on different cell types.

| Dataset | RMSE | Avg. Accuracy | R2 value | Pearson's Correlation |
| --- | --- | --- | --- | --- |
| H358 | 0.00406 | 91.87 | 0.8815 | 0.9389 |
| Broad u2os | 0.01278 | 87.857 | 0.9054 | 0.9515 |
| KP/KL | 0.00536 | 90.543 | 0.9662 | 0.9829 |
| Overall Average |  | 90.09 | 0.9177 | 0.9576 |

## Discussion

In this study, we present *micronuclAI*, a novel framework that harnesses the power of deep-learning technology and computer vision techniques for CIN assessment in cancer cell lines *in situ*. By focusing on MN quantification, *micronuclAI* offers a reliable proxy for assessing CIN at scale, providing a standardized and efficient solution.

Prior efforts recognized the importance of automating MN quantification ^39,40^, typically through combinations of traditional computer vision, machine learning or deep learning approaches with their own advantages and limitations as shown in Fig. 1 and Table 1. A large number of previous works have used the Cytokinesis-block MicroNucleus (CBMN) assay following the protocols defined by the HUMN (HUman MicroNucleus) project ^41^ to quantify MN within binucleated lymphocytes. Briefly, the CBMN assay is a standardized method for genotoxicity studies where biomarkers of CIN are scored in binucleated cells after cytokinesis is stopped by addition of cytochalasin B ^39,42^. Other exemplary methods automate MN quantification in various other cell/tissue types after genotoxic exposures, citing no difference in results between the *in vitro* MN tests with or without the use of cytokinesis blockers like cytochalasin B ^43,44^. Meanwhile, other methods have proposed quantification of nuclear budding MN specifically ^28^, or focus on a combination of MN and toroidal nuclei ^31^ to quantify CIN. Importantly, to our knowledge there have been no published methods for automatic quantification of MN of varying morphology, localization, across species and lineages in cancer cell lines. In addition, most methods evaluate their performance on a limited dataset plagued by inter-observer variability. Therefore, existing methods face certain limitations such as i) quantification specifically of micronuclei alone within binucleated lymphocytes, ii) quantification of NBUDs alone, iii) requirement of cellular/cytoplasmic staining in addition to nuclear staining, iv) lack of adequate evaluation, and v) are not open-source or easily available. In contrast, micronuclAI performs well across multiple biological, perturbational, and technical conditions, and achieves human-level performance at a fraction of the time, taking approximately 10 seconds (on a MacBook Pro M1) to score 3000 cells compared to the ∼120 minutes for manual scoring. Its robustness extends to handling multiple nuclear staining, as demonstrated with DAPI and Hoechst datasets. Furthermore, micronuclAI proves its scale invariance by excelling under varying microscopy conditions, including 10X, and 100X objective datasets.

Like many AI systems, micronuclAI has limitations. It cannot explicitly communicate the reasoning behind its predictions, but its value lies in its consistent and scalable detection of micronuclei as a reliable indicator of CIN. We partly address this issue by using saliency maps to highlight the regions in the image that contributed the most to drive the prediction. Another limitation is the reliance on a nuclear segmentation mask. Although precise identification of the nuclear boundaries is not required and the method is robust to multiple segmentation methods, instances of extensive mis-, over-, and under-segmentation may affect the predicting capabilities of the framework so the segmentation quality cannot be fully omitted. Additionally, instances of the same MN on two different isolated nuclei images might be present, leading to a potential double counting of the same structures within two different patches. Such instances occur on areas of the image with high nuclear/cell density that might yield misleading or inflated results. Although rare, these instances can be minimized by following two approaches: 1) changing the FOV used for the nuclei isolation step and, 2) addressing the experimental design before imaging. To quantify the frequency of these occurrences, we calculated the Intersection over Union (IoU) between all the patches in an image. 0.00168% of the pairs had an IoU >= 0.5 signifying that only a negligible amount of overlap is present in the dataset, hence we can safely disregard the concern for double counting - especially when also controlling for cell density. We have directed efforts to mitigate and understand these challenges.

In summary, micronuclAI is a robust and scalable tool for quantifying CIN, and will be a critical tool for understanding CIN biology and its role in tumor biology and treatment responses.

## Methods

### Cell culture

A375 and NCI-H358 were obtained from the American Type Culture Collection (ATCC), whereas HCC44 was obtained from the Leibniz Institute DSMZ. KP (*Kras^G12D^Tp53*^-/-^) and KL (*Kras^G12D^Lkb1*^-/-^) were kindly gifted from Kwok-Kin Wong. NCI-H358, HCC44, KP, and KL were maintained in RPMI culture media supplemented with 10% FBS and 1% penicillin/streptomycin, whereas A375 were cultured in DMEM, supplemented with 10% FBS and 1% penicillin/streptomycin.

### Data Acquisition

Cells were seeded at a density of 1500 cells/well in opaque bottom 96-well tissue culture plates (Corning). Once confluency reached 70-80% in the well, cells were fixed in 4% paraformaldehyde for 15 minutes. After washing with PBS, cells were incubated with Hoescht (Thermo Fisher) diluted in Odyssey Blocking Buffer (1:10000). Fluorescent images were obtained on the Zeiss Celldiscoverer 7 using the PlanApochromat 20X/0.7 objective, 0.5X magnification changer, and Axiocam 506. Whole-well images were stitched and exported using Zen 3.1 Software resulting in a final resolution of 8829 x 9072 pixels.

### Segmentation

We extracted only the nuclear channel from raw images as ome.tif files using AICSImageIO (v 4.11.0) ^45^. We performed nuclear segmentation on the resulting ome.tif images using: DeepCell Mesmer Whole-Cell (v 0.4.1), DeepCell Mesmer Nuclear (v 0.4.1), DeepCell Nuclear Segmentation (v 0.4.1) ^34^, Stardist (v 0.8.3) ^36^ and Cellpose (v 2.2.2) ^35^ on a high-performance computational cluster, bwHPC helix. DeepCell Mesmer and DeepCell Nuclear are deep-learning models used for cell segmentation and nuclear segmentation. DeepCell models are trained on a large number of images to achieve human level performance in the segmentation task. While DeepCell mesmer is trained on histological images DeepCell Nuclear is optimized for segmentation on images derived from cell cultures. Cellpose is a human-in-the-loop generalist model for cell segmentation, providing some pre-trained models. In this case we used the ‘nuclei’ model to perform nuclei segmentation. Stardist is a deep-learning based object detection with Star-convex shapes algorithm used for 2D and 3D object detection and segmentation in microscopy. Training data was generated using Stardist generated segmentation masks and the corresponding nuclear .ome.tif files.

### Nuclei Isolation & Preprocessing

Using the segmentation mask, we obtained the coordinates that circumscribe each of the segmented nuclei. We then use these coordinates to extract isolated nuclei (IN) patches from the nuclear images. To homogenize the apparent cell sizes of each single nuclei we calculate a scaling factor for each one. The factor is calculated with the following formula:

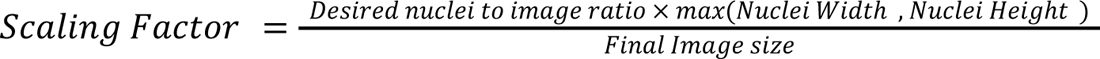

In this study, we use a *nucleus to image ratio* of 0.65; meaning each isolated nuclei occupies sixty five percent of the final image size. We increase the field of view (FOV) around each isolated nucleus by expanding the circumscribed bounding box (20 pixels in the case of the training data). To remove segmentation errors from the pipeline, we remove the bounding boxes falling into the first five percentile by area. The remaining isolated nuclei images are then resized and cropped to the center to a final resolution of 256 x 256 pixels. Lastly, we correct the brightness intensity in each image with brightness normalization. To facilitate this process, we developed the *mask2bbox* Python library, which is specifically designed for generating, handling, and visualizing bounding boxes from a segmentation mask image. Mask2bbox is available as a python package via the Python Package Index (PyPI), Bioconda, and as Docker and Singularity containers.

### Data Labeling

A total of 84,286 isolated nuclei patches from the A375 cell line images at 10x were labeled by experts using a PySimpleGUI-based (v 4.60.4) ^46^ labeling tool. This tool is available to use as a PyPI and Conda package. To label the micronuclei, we adapted the protocol established by the HUMN project ^41^ for human lymphocytes, and the scoring criteria in oral exfoliated cells from ^47^ with some adjustments given the cell lines used and downstream application.

A micronuclei is counted if it fulfills the following criteria:

a. Rounded and smooth perimeter suggestive of a membrane if separate from the primary nuclei.
b. Less than one-third of the diameter of the associated nucleus, but large enough to discern the shape.
c. Staining intensity similar to that of the primary nucleus.
d. Texture similar to that of the primary nucleus.
e. Same focal plane as the primary nucleus.

A NBUD is counted if it fulfills the above criteria for micronuclei and appears to be touching or budding out of the primary nuclei but is clearly distinguishable from/within the primary nuclear boundary.

Additionally, apoptotic and mitotic cells were identified with the following criteria and labeled as 0 CIN count.

a. Apoptotic cells: Single-cell patches with a large number of fragmented objects and no definitive primary nuclei were considered as apoptotic nuclei.
b. Mitotic cells: Single-cell patches with very bright double nuclei were considered as mitotic nuclei.

### Model Training

We used the labeled isolated nuclei images to train a convolutional neural network (CNN) to quantify the number of micronuclei associated with each primary nucleus. The quantification is achieved using a model based on the *EfficientNet* V1 architecture. The model architecture (Fig. 3A), is composed of a fully connected CNN, followed by a channel-wise convolution component as a feature extractor, and a fully connected layer for the final prediction. We employ versions B0 to B7 of *EfficientNet* V1, which are models of increasing complexity, as the backbone feature extractors. The models were implemented in *Pytorch* (v 1.13.0) ^48^ using the *Pytorch lightning* framework (v 1.8.0) ^48^

The models were trained using the mean squared error (MSE) as the loss function and Adaptive moment estimation (ADAM) as the gradient descent optimizer. The initial learning rate was set to 10e-3 and decayed using ReduceLRonPlateu (with a factor of 0.2 and a patience of 10 epochs) with an early stop criteria to a maximum of 300 epochs. For regularization, a dropout value of 0,2 was added to all fully connected (FC) layers. All EfficientNet (B0-B7) models were trained both with random initial weights and with weights of models pre-trained on ImageNet. Data augmentations were applied at random during training which include random vertical flip, random horizontal flip, random rotation ± 0 - 30 degrees, randomly applied gaussian blur (p=0.3, kernel_size = (3, 3)) and data normalization [-1, 1]. The mini-batch size was set to 64 for all models.

A 90:10 training:validation split was used for all the models. We evaluated and selected the final weights for each model based on the lowest root mean squared error (RMSE) value on the validation set. Furthermore, the models were compared to each other based on standard metrics, such as F1-score, precision-recall curves and AUPRC. Prediction values were rounded to the nearest integer for comparison with the training labels.

### Manual data annotation

In addition to the manual labeling performed to generate the training data, we manually counted nuclei and MN on the cell lines used for validation of the complete pipeline. We used FIJI’s (v 2.14.0 / 1.54f) ^49^ multi-point tool to count all nuclei present in an image while separately keeping track of all the MN and NBUDs. These values were then used to estimate the degree of CIN by calculating the CIN score, as a ratio between the total number of MN and total number of nuclei for each image. A Monte Carlo approach was used to estimate the CIN score in images where the number of cells was too large for manual labeling. This Monte Carlo estimation consisted of randomly selecting 3 FOV from each image and assessing MNs and NBUDs for approximately 800 to 1000 nuclei per FOV. A summary of all the available data is present in Supplementary Table 1.

### Evaluation

#### Evaluation of the CNN model

We evaluated the CNN model on the hold-out test dataset within the initial isolated nuclei images. The model prediction values for the CIN count associated with each isolated nuclei were rounded to the closest integer value. The model performance was then assessed with standard classification metric, F1-score and Precision-Recall curve using the scikit-learn implementation in Python (v 1.0.2) ^50^. We also generated attention maps with Class Activation Maps (CAM) ^51^ as a part of our attempt at generating explainable AI (XAI). Attention maps provide a visual representation of how the neural network processed the information helping us understand and confirm the model’s decisions.

#### Evaluation of the pipeline on the H358 cell-line

We evaluated the performance of the pipeline on images taken from the H358 cell-line, a primary bronchoalveolar carcinoma of the lung, a non-small-cell lung cancer. The dataset contains 17 DAPI-stained images, 10k x 10k pixels imaged at 10x containing approximately 20,000 nuclei in total. This set of images consisted of 3 different genetic variations that alter the CIN level: H358, H358-GFP, H358-dnMCAK in order of CIN level from low to high. The evaluation was done on the basis of CIN score for each image compared to manual expert quantifications. R^2^ and Pearson’s correlation were calculated as a summary statistic.

#### Evaluation of the pipeline on an external dataset imaged at 100x

To test the scale invariability of the method, we evaluated the performance of micronuclAI with an external image dataset BBBC039v1 ^38^, available from the Broad Bioimage Benchmark Collection ^52^. The dataset contained Hoechst-stained human U2OS cells with 200 fields of view, 520×296 pixels imaged at a 100x zoom. Evaluation was done on a set of 20 random images from this dataset containing approximately 2400 nuclei by comparing the manual and micronuclAI estimated CIN score using both R^2^ and Pearson correlation as summary statistics.

#### Evaluation of the pipeline on murine cell lines

We further tested micronuclAI on mice derived KP/KL cell lines: KP (*Kras^G12C^Tp53*^-/-^) and KL (*Kras^G12C^Lkb1*^-/-^). 5 KP and 5 KL images were manually annotated and compared to micronuclAI predictions. Given the large number of cells images (∼20 K per image) we used the Monte Carlo estimation counting strategy for this task.

#### Implementation / Deployment

The complete micronuclAI pipeline is available as a Command Line Interface (CLI) through Github and a simple Graphical User Interface (GUI) through Streamlit ^53^. Inference of micronuclei can be achieved in small to medium sized example images that can be uploaded to the streamlit app. Image data is processed within a virtual machine (VM) on Heicloud ^54^, a local Cloud infrastructure provided by University Computing Center Heidelberg, and images are immediately deleted after micronuclei inference. Once micronuclei are inferred, results predictions as well as several plots describing the results are generated and presented to the user within the streamlit app. No data is kept on the server after the user disconnects from the app.

## Data Availability

Training data and labels

https://www.synapse.org/#!Synapse:syn54780485/

Broad Bioimage Benchmark Collection

https://bbbc.broadinstitute.org/

## Code Availability

Labeling Tool

Github: https://github.com/SchapiroLabor/micronuclAI_labeling

Mask2BBox

Github: https://github.com/SchapiroLabor/mask2bbox

micronuclAI

Github: https://github.com/SchapiroLabor/micronuclAI

Streamlit App

Github: https://github.com/SchapiroLabor/micronuclAI/streamlit

## Acknowledgements

The authors acknowledge support by the state of Baden-Württemberg through bwHPC and the German Research Foundation (DFG) through grant INST 35/1597-1 FUGG. A.H. was supported through state funds approved by the State Parliament of Baden-Württemberg for the Innovation Campus Health + Life Science Alliance Heidelberg Mannheim. D.S., M.A.I.A., N.S. and F.W. was supported by the German Federal Ministry of Education and Research (BMBF 01ZZ2004). F.W. was supported by a Walter-Benjamin position from the Deutsche Forschungsgemeinschaft (DFG). B.I. is supported by National Institute of Health grants, R37CA258829, R01CA280414, R01CA266446, U54CA274506, and additionally by the Pershing Square Sohn Cancer Research Alliance Award, the Burroughs Wellcome Fund Career Award for Medical Scientists; a Tara Miller Melanoma Research Alliance Young Investigator Award; the Louis V. Gerstner, Jr. Scholars Program; and the V Foundation Scholars Award. L.A.C is supported by NIH/NCI grant F30CA281104-01 and, along with L.C., MSTP Training Grant T32GM007367. This work is supported by the Health + Life Science Alliance Heidelberg Mannheim and received state funds approved by the State Parliament of Baden-Württemberg. (“AI Health Innovation Cluster” and “MULTI-SPACE”). We thank our administrative and project management team: Erika Schulz, Lydia Roeder and Bettina Haase. We would also like to thank Ricardo Omar Ramirez Flores, Jovan Tanevski, Victor Perez, Krešimir Beštak, and Chiara Schiller for the helpful discussions and Cristina-Ruxandra Burghelea for her help in proofreading the manuscript.

## Author Contributions

M.A.I.A.: Methodology, Interpretation, Writing, Figures L.A.C.: Data Acquisition, Methodology, Interpretation, Writing A.H.: Methodology, Interpretation, Writing F.W: Streamlit app. N.S.: Methodology L.C.: Data Acquisition S.S.: Data acquisition J.C.M.: Data acquisition, Methodology B.I.: Conceptualization, Interpretation, Writing, Supervision, Funding D.S: Conceptualization, Interpretation, Writing, Supervision, Funding

## Supplementary Tables

**Supplementary Table 1:** List of annotated datasets.

| Dataset | Cell line | Modifications | No. of Images | No. of Nuclei | Objective | Staining |
| --- | --- | --- | --- | --- | --- | --- |
| A375 | Malignant Melanoma | control/ Kif2a<br>MCAK/dnMC<br>AK | 23 | 84,286 | 10 | DAPI |
| H358 | Human Non-small cell lung cancer | control/ GFP/<br>dnMCAK | 17 | 19,954 | 10 | DAPI |
| Broad<br>BBBC039v1 | Human osteosarcoma (u2os) | none | 20 | 1,999 | 100 | Hoechst |
| KP/KL | Mouse Non-small cell lung cancer | none | 12 | 9,237 | 10 | DAPI |
| Total |  |  | 72 | 115,476 |  |  |

## Supplementary figures

**Supplementary Figure 1:**
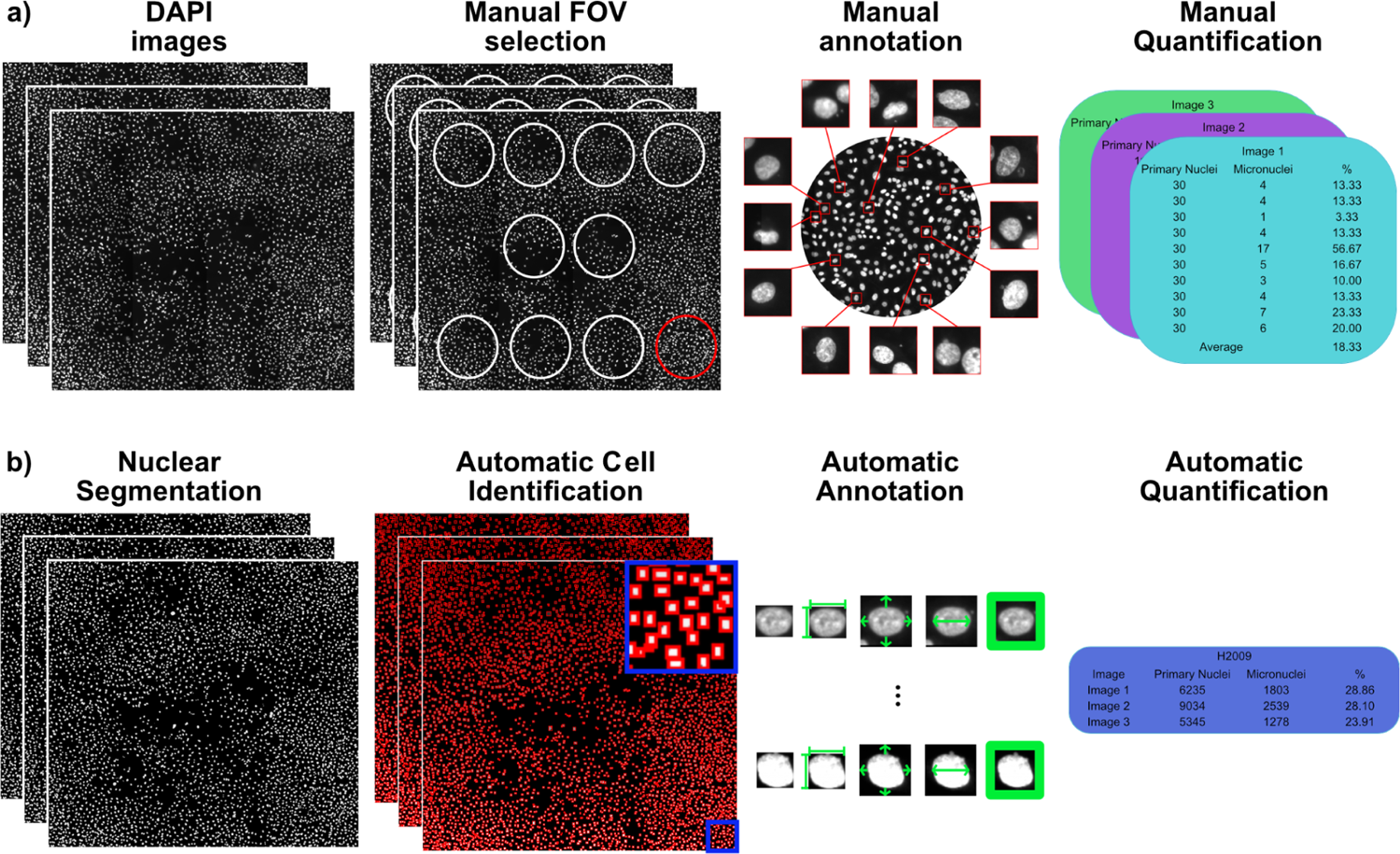
Detailed workflow of the manual quantification and micronuclAI. **a-b,** Details of the manual workflow **(a)** for micronuclei quantification and the proposed automatic quantification workflow with micronuclAI **(b)**.

**Supplementary Figure 2:**
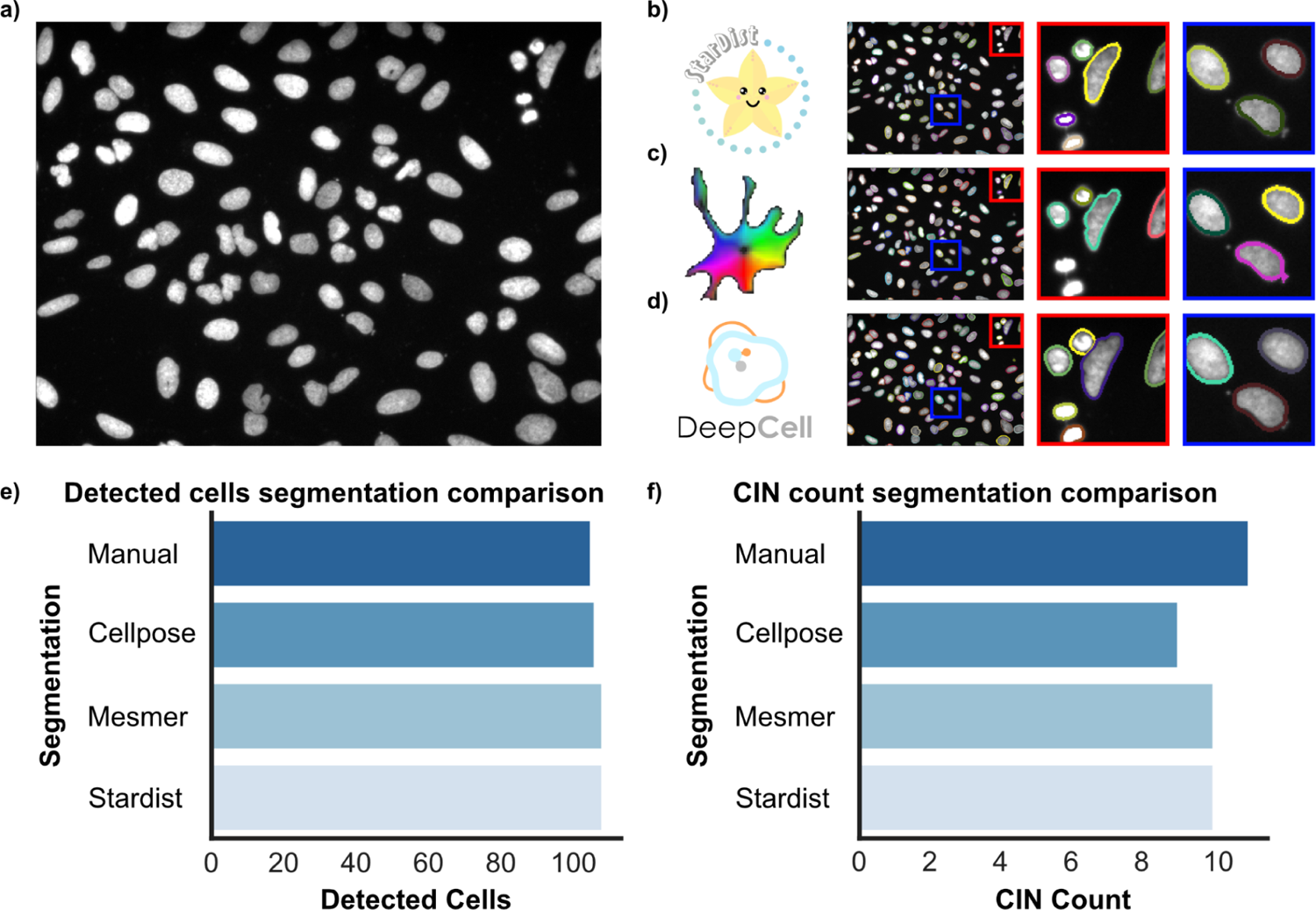
Example of various segmentation methods on CIN predictions. **a-d,** Example of Hoechst stained image **(a)** and the segmentation overlays with zoom-in regions for three segmentation methods: Stardist **(b)**, Cellpose **(c)** and DeepCell **(d)** nuclear segmentation. **e-f,** Comparison of the number of detected cells **(e)** and CIN counts **(f)** across manual counting and the segmentation methods.

